# Metabolome-Wide Mendelian Randomization Analysis of Emotional and Behavioral Responses to Traumatic Stress

**DOI:** 10.1101/545442

**Authors:** Carolina Muniz Carvalho, Frank R. Wendt, Dan J. Stein, Murray B. Stein, Joel Gelernter, Sintia I. Belangero, Renato Polimanti

## Abstract

Trauma exposure is an important risk factor for several psychiatric disorders; however, the mechanisms that underlie emotional and behavioral responses to traumatic stress are unclear. To understand these mechanisms, this study investigated the genetic overlap and causal relationship between blood metabolites and traits related to trauma response using genome-wide data. Five traits related to trauma response “in the past month” ascertained in the UK Biobank (52 816<N<117 900 individuals) were considered: i) “Avoided activities or situations because of previous stressful experience” (*Avoidance*); ii) “Felt distant from other people” (*Distant*); iii) “Felt irritable or had angry outbursts” (*Irritable*); iv) “Felt very upset when reminded of stressful experience” (*Upset*); v) “Repeated disturbing thoughts of stressful experience” (*Repeated Thoughts*). These were investigated with respect to 52 metabolites assessed using nuclear magnetic resonance metabolomics in a previous genome-wide association study (up to 24,925 individuals of European descent). Applying linkage disequilibrium score regression (LDSC), polygenic risk scoring (PRS), and Mendelian randomization (MR), we observed that 14 metabolites were significantly correlated with trauma response traits (p<0.05); PRS of 4 metabolites (citrate (CIT); glycoprotein acetyls (GP); concentration of large very-low-density lipoproteins (VLDL) particles (LVLDLP); total cholesterol in medium particles of VLDL (MVLDLC)) were associated with traits related to trauma response (false discovery rate Q<10%). These associations were partially due to causal relationships (CIT→Upset β=-0.058, p=9.1×10^−4^; GP→Avoidance β=0.008, p=0.003; LVLDLP→Distant β=0.008, p=0.022; MVLDLC→Avoidance β=0.019, p=3×10^−4^). No reverse associations were observed. In conclusion, the genetics of certain blood-metabolites are potentially implicated in the response to traumatic experience.

## Introduction

Psychiatric disorders are comprised of multifactorial traits characterized by the interaction between multiple genes and environmental factors (1). Early child adversity and later exposure to traumatic events, for example, are key risk factors for multiple mental health disorders such as posttraumatic stress disorder (PTSD) (2) and depression (3). Approximately 60–90% of individuals experience at least one traumatic or violent event during their lifetimes, which can trigger a stress response (4, 5), leading to transient psychiatric symptoms and, in a relatively small proportion of trauma-exposed individuals, psychiatric disorders (5, 6). This susceptibility may be attributed to different factors including genetic and epigenetic variation, previous exposure to traumatic events, and neurobiological alterations (7, 8).

The stress response after traumatic events involves a range of biological and behavioral systems, and leads to alterations in neural, endocrine, and immune activity. These alterations may in turn increase the risk of developing diseases (9–12). Prior studies have focused on investigation of genomics, transcriptomics, and epigenomics (13–15) aiming to understand the role of trauma in the etiology of psychiatric disorders and to identify genetic biomarkers for these conditions. Genome-wide association studies (GWAS) have found evidence of genetic associations with numerous psychiatric disorders (15–20), but evidence for the influence of genetic factors on the biological processes underlying the trauma response is still preliminary. Recent GWAS of PTSD suggested that more studies are necessary to identify robust genetic variants associated with PTSD due to complexity and heterogeneity of this phenotype (21).

Evaluation of metabolites is a new approach to identify disease pathophysiological components. Metabolites are a group of end-products of cellular regulatory processes that potentially provide insight into those processes and are capable of informing the complex relationship between genotype and phenotype (22, 23). Psychiatric disorders, cognition, and dementia have been associated with alterations in circulating metabolites (23–26). Despite this, the causal relationship between metabolite levels and biological mechanisms underlying these phenotypes remains unclear.

Mendelian randomization (MR) is a method that relies on genetic data to infer causal relationships, for example, between an exposure (e.g., a metabolite) and an outcome (e.g., psychiatric disorder) using genetic variants as instrumental variables (27–31). There are few studies evaluating the role of metabolites on phenotypes resulting from exposure to trauma and no study to date has examined the putative causality between these factors using the MR approach. Therefore, the aim of this study was to investigate the genetic correlation between trauma-related traits and metabolites, as well as their causal relationship.

## Materials and Methods

### Study Design and Data Sources

We conducted two sample MR (32) to estimate the causal relationship between genetic variants associated with metabolite levels and traits related to trauma response using publicly available summary data.

GWAS summary data pertaining to phenotypic traits included in the category “traumatic events” (Field ID: 145) were obtained from the UK Biobank (UKB) (33, 34). UKB is an open-access international health research which provides genetic information about thousands of individuals (aged 40–69 years) (33, 34). For characterization of mental health phenotypes, all participants (including females and males, N=157 366) provide a self-report about lifetime symptoms of mental disorders answering an online mental health questionnaire (MHQ), which includes trauma measures that were previously validated (35).

Among the traits available in this UKB category, we selected those related to trauma response based on recent response to stressful past experiences from the UKB online "Thoughts and Feelings" MHQ about: “Avoided activities or situations because of previous stressful experience in past month” (*Avoidance*; UKB Field ID: 20495; N=117 868), “Felt distant from other people in past month” (*Distant*; UKB Field ID: 20496; N=52 822), “Felt irritable or had angry outbursts in past month” (*Irritable*; UKB Field ID: 20494; N=52 816), “Felt very upset when reminded of stressful experience in past month” (*Upset*; UKB Field ID: 20498; N=117 893), and “Repeated disturbing thoughts of stressful experience in past month” (*Repeated Thoughts*; UKB Field ID: 20497; N=117 900) and the answers options for these questions were: “Not at all”, “A little bit”, “Moderately”, “Quite a bit”, “Extremely”. Supplemental Figure S1 shows the distribution of UKB participants’ answers to the trauma-response questions. These responses can also be thought of as common symptoms of PTSD, i.e., re-experiencing, avoidance, and hyperarousal (36, 37).

To investigate the metabolic traits, we used the GWAS data published by MAGNETIC NMR (nuclear magnetic resonance) study, which investigated 123 circulating metabolites quantified by nuclear magnetic resonance metabolomics in up to 24 925 individuals of European descent (38). The characteristics of each GWAS used in this study are provided in Table 1. For more information about the quality control and GWAS methods applied in relation to trauma-related traits (UKB) and metabolites see, respectively, https://github.com/Nealelab/UK_Biobank_GWAS/tree/master/imputed-v2-gwas and MAGNETIC NMR GWAS summary statistics (http://www.computationalmedicine.fi/data#NMR_GWAS) (38).

**Table 1:** Characteristics of cohorts evaluated

| Trait | Abbreviation | Sample Size |
| --- | --- | --- |
| Felt irritable or had angry outbursts in past month (UKB Field ID: 20494) | Irritable | 52 816 |
| Avoided activities or situations because of previous stressful experience in past month (UKB Field ID: 20495) | Avoidance | 117 868 |
| Felt distant from other people in past month (UKB Field ID: 20496) | Distant | 52 822 |
| Repeated disturbing thoughts of stressful experience in past month (UKB Field ID: 20497) | Repeated Thoughts | 117 900 |
| Felt very upset when reminded of stressful experience in past month (UKB Field ID: 20498) | Upset | 117 893 |
| Metabolites | -- | 24 925 |

UKB: UK Biobank

### Heritability and Genetic Correlation

Linkage disequilibrium score regression (LDSC) was used to calculate metabolite heritability estimates and to test the genetic correlation between trauma-related traits and metabolites. Also, we calculated the genetic correlation among the five traits related to trauma response to verify how these traits are correlated with each other.

In addition, we derived the heritability z-score for all traits, which is defined as the heritability estimate produced by LDSC regression divided by its standard error. The value of z-score may be affected by sample size, SNP-based heritability and the proportion of causal variants (39). In other words, the heritability z-score can capture information about the genetic basis of a trait. In our analyses, we considered heritability z-score > 3 as suitable to conduct the genetic correlation analysis.

Summary statistics were formatted according to the pipeline described by Bulik-Sullivan et al. (40, 41). We used the HapMap 3 reference panel (42) and pre-computed LD scores based on the 1000 Genomes Project data (43) for European ancestry (available at https://github.com/bulik/ldsc) to estimate the genetic correlation between traits.

### Polygenic Risk Score and Definition of the Genetic Instruments

We conducted a polygenic risk score (PRS) analysis to define the genetic instruments for MR using PRSice v1.25 software (44). PRS was calculated using the following parameters: clumping with an LD cutoff of R^2^ = 0.001 within a 10 000-kb window and removal of the major histocompatibility complex (MHC) region of the genome because of its complex LD structure. European samples from the 1000 Genomes Project were used as the LD reference panel.

PRS analysis was conducted using GWAS summary data for both base and target datasets (i.e., traits related to trauma response and blood metabolites). The p-values were adjusted for multiple testing correction using the false discovery rate (FDR) and we considered Q < 10% as the significance threshold.

### Mendelian Randomization

To assess causal relationship between the phenotypes investigated in this study, we selected single nucleotide polymorphisms (SNPs) associated with the risk factor according to the best-fit PRS estimates for each metabolite surviving multiple testing correction (FDR Q < 10%).

To estimate the causal relationship between trauma-related traits and metabolites, we used different MR methods available in the TwoSampleMR R package (32): random-effects inverse variance weighted (IVW) (32), MR-Egger (45), weighted median (46), simple mode (47) and weighted mode (46).

We conducted sensitivity analyses to estimate the heterogeneity of the data using the IVW and MR-Egger heterogeneity tests to exclude the presence of possible biases in the MR estimates. Additionally, we conducted a reverse analysis with respect to these phenotypes to verify whether traits related to trauma response have a causal effect on the metabolite levels. Furthermore, we performed MR-Egger regression intercept and a MR-PRESSO (Pleiotropy RESidual Sum and Outlier) global test to estimate pleiotropy (48). We used a leave-one-out analysis to identify potential outliers in the genetic instruments indicative of significant pleiotropy or heterogeneity.

### Enrichment analysis

Based on the MR results, we conducted Gene Ontology (GO) enrichment analysis to investigate possible biological, molecular, or cellular processes associated with metabolite levels using eSNPO (eQTL based SNP Ontology) (49). We performed the enrichment analysis for eQTLs present in two tissues: blood and brain. FDR multiple testing correction was applied to adjust the results of these analyses considering Q < 5% as the significance threshold. The GO enrichment results were analyzed using REVIGO (http://revigo.irb.hr/) to generate a graph-based visualization (50), which was done considering a similarity of 0.7 between GO terms, UniProt as the reference database, and the Jian and Conrath method as the semantic similarity measure.

## Results

### Genetic Correlation

SNP heritability was calculated with respect to 123 blood metabolites (Supplemental Table S1). Considering those metabolites with strong heritability estimates (z score > 3), we observed 52 traits with heritability ranging from 6.1% (valine) to 16.6% (double bonds in fatty acids). We observed 17 genetic correlations (p < 0.05) with respect to these 52 metabolites and the five UKB traits related to trauma response (Figure 1). The strongest genetic correlation was observed between glycoprotein acetyls (GP) and *Upset* (rg = 0.318, p = 0.003). The same metabolite was also genetically correlated with *Avoidance* and *Repeated Thoughts* (rg = 0.349, p = 0.004; rg = 0.221, p = 0.034; respectively). This is due to the fact that the traits related to trauma response investigated showed a high genetic correlation among each other (Supplemental Table S2).

**Figure 1:**
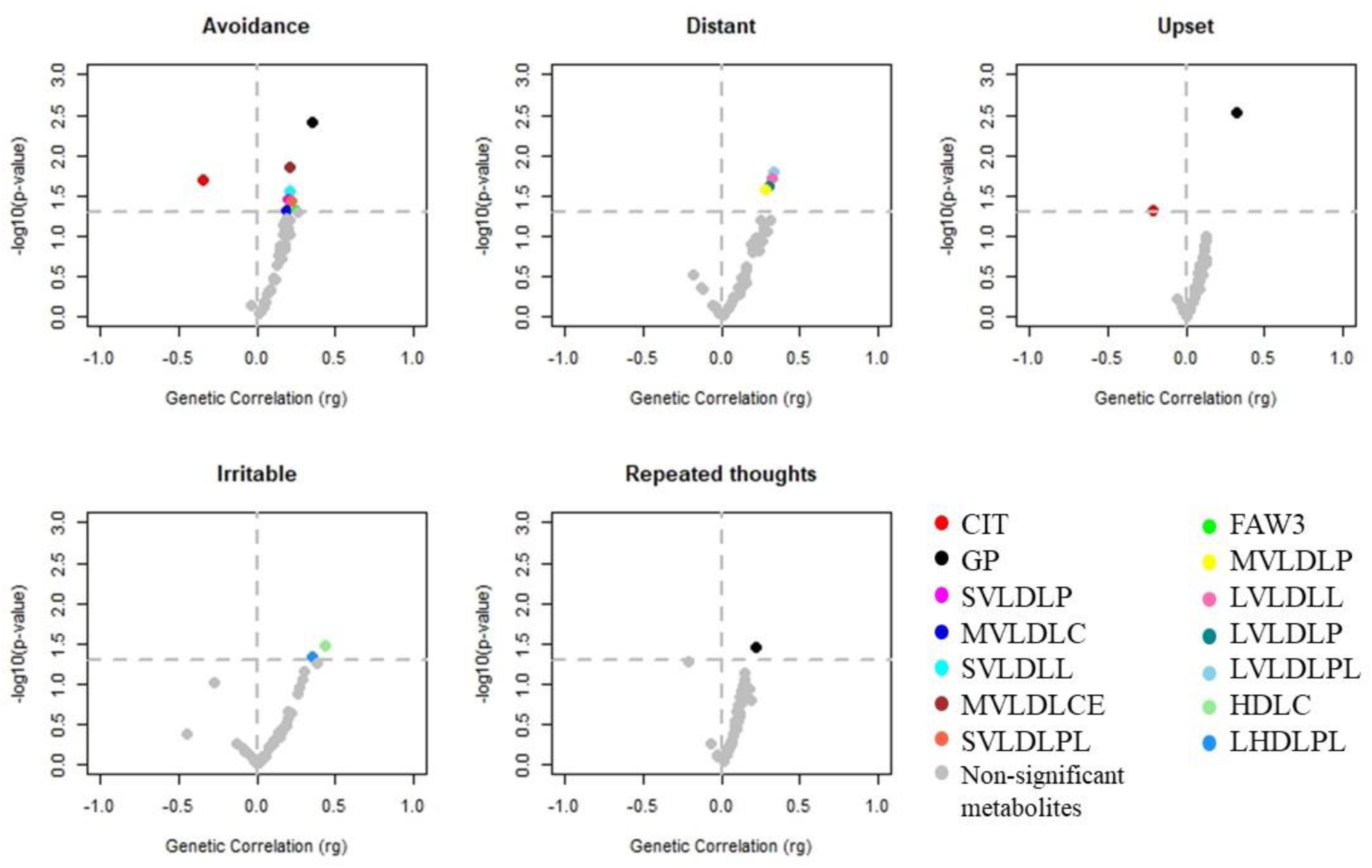
Genetic correlation results between metabolites and traits related to trauma response. *CIT*: Citrate. *FAW3*: Omega-3 fatty acids. *GP*: Glycoproteins acetyls. *HDLC*: Total cholesterol in HDL (High-density lipoprotein). *LHDLPL*: Phospholipids in large HDL. *LVLDLL*: Total lipids in large VLDL (Very-low-density lipoprotein). *LVLDLP*: Concentration of large VLDL particles. *LVLDLPL*: Phospholipids in large VLDL. *MVLDLC*: Total cholesterol in medium VLDL. *MVLDLCE*: Cholesterol esters in medium VLDL. *MVLDLP*: Concentration of medium VLDL particles. *SVLDLL*: Total lipids in small VLDL. *SVLDLP*: Concentration of small VLDL particles. *SVLDLPL*: Phospholipids in small VLDL.

### Polygenic Risk Score and Definition of the Genetic Instruments

Given the 17 genetic correlations observed, we conducted a PRS analysis to investigate these genetic overlaps further and to determine genetic instruments to use in the MR analysis. After FDR 10% correction for the number of PRS thresholds tested, 5 associations remain significant (Table 2). GP PRS was associated with *Avoidance* and *Repeated Thoughts* (FDR Q = 0.06 and Q = 0.057, respectively). Citrate PRS was associated with *Upset* phenotype (FDR Q = 0.006). The two remaining associations were observed with respect to two subclasses of very low-density lipoproteins (VLDL): concentration of large very-low-density particles (LVLDLP) was associated with *Distant* (FDR Q = 0.018) and the total cholesterol in medium of very-low-density lipoprotein (MVLDLC) was associated with *Avoidance* (FDR Q = 0.055). No reverse association between PRS of trauma-response phenotype and these blood metabolites was found (FDR Q > 0.1).

**Table 2:**
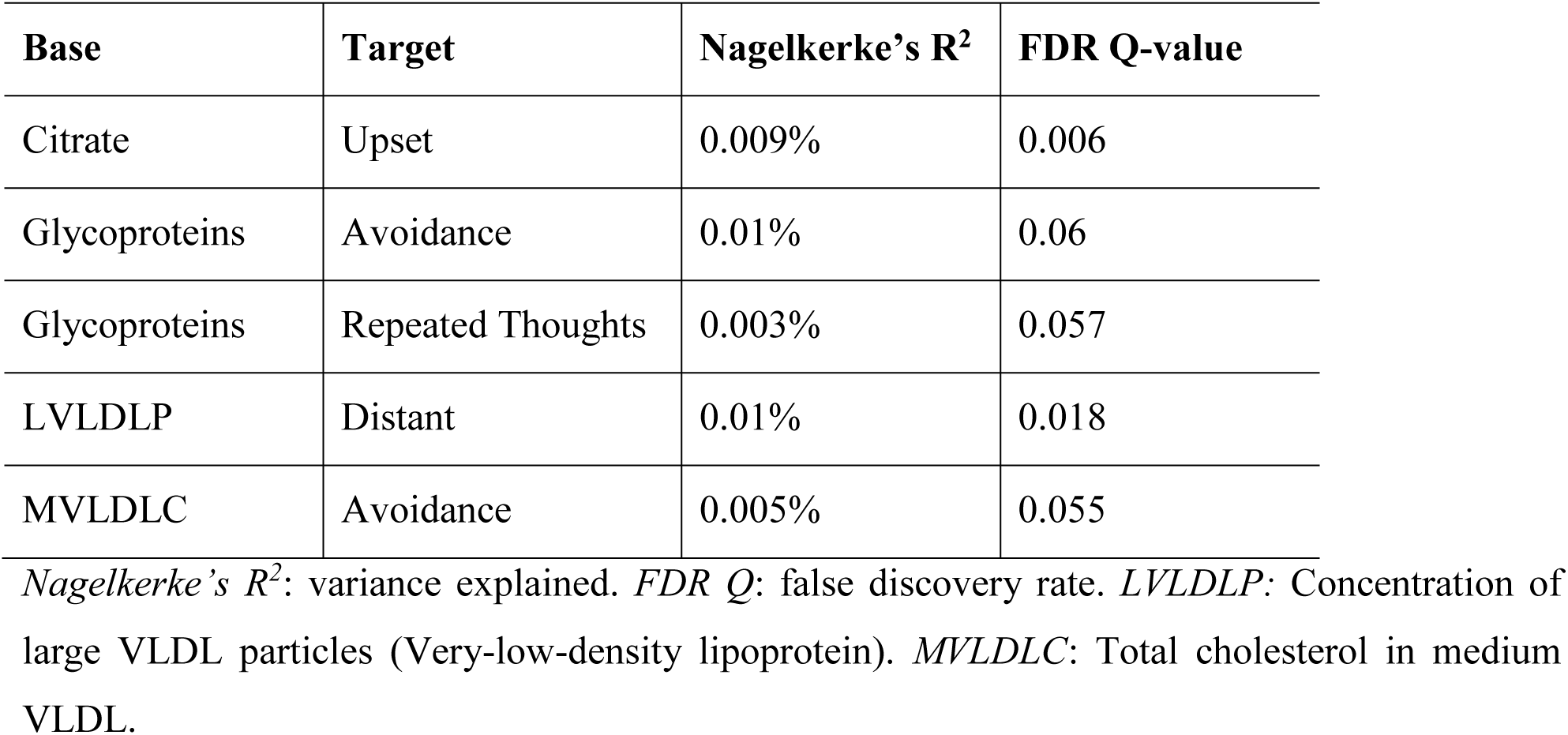
PRS results

### Mendelian Randomization

Based on the PRS associations, we assessed the possible causal relationships using MR methods (Figure 2a, Supplemental Table S3). We found a negative causal effect of citrate levels on the *Upset* trauma response trait (IVW β = −0.058, p = 9.1×10^−4^). This estimate was consistent across multiple MR methods (Supplemental Table S3) and no heterogeneity or pleiotropy was observed in this analysis (Supplemental Tables S4 and S5).

**Figure 2:**
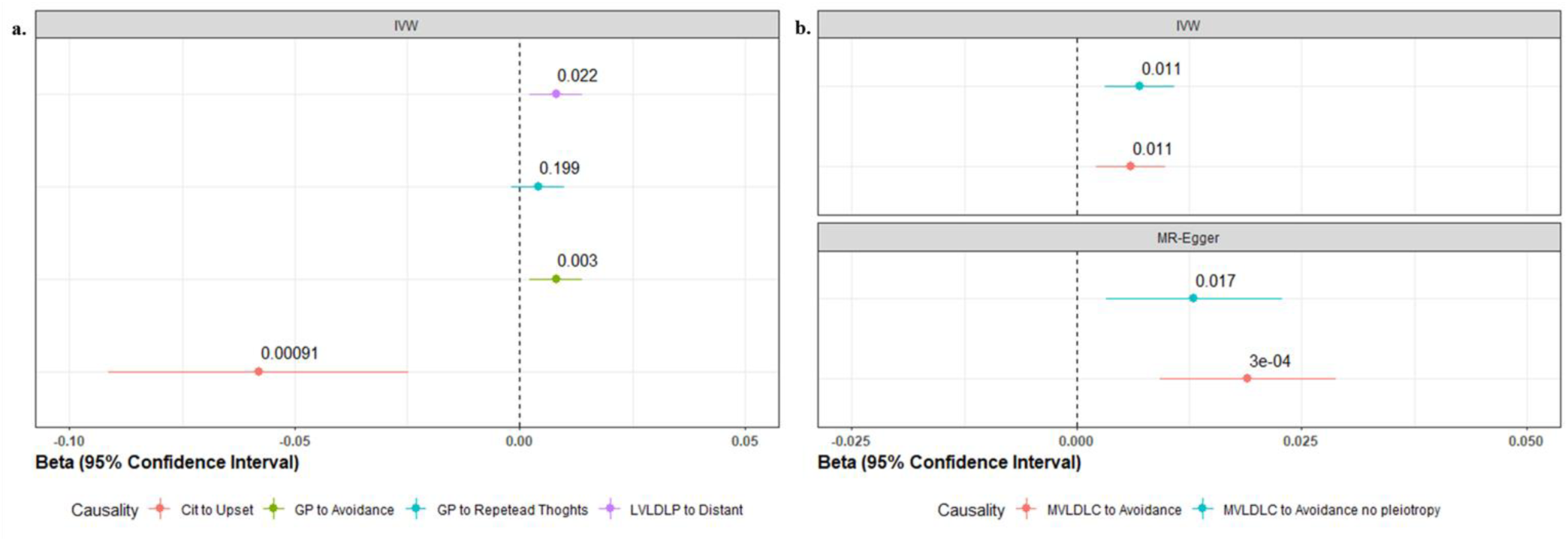
Mendelian randomization (MR) summary. **a.** Causal effects (beta) and p-values of IVW method for the relationship of the SNP effects on the metabolites against the SNP effects on the traits related to trauma response. **b.** Causal effects (beta) and p-values of IVW and MR-Egger for the relationship of the MVLDLC on *Avoidance* considering variants with pleiotropy (in red) and no pleiotropy (in cyan). *Cit:* Citrate. *GP:* Glycoproteins acetyls. *LVLDLP*: Concentration of large VLDL particles (Very-low-density lipoprotein). *MVLDLC*: Total cholesterol in medium VLDL. *IVW:* inverse variance weighted.

We verified that the GP levels appear to have a small causal effect on *Avoidance* (IVW β: 0.008, p = 0.003), but did not have an effect on *Repeated Thoughts* (p-value > 0.05). No evidence of bias due to pleiotropy and heterogeneity was observed in the GP genetic instrument (Supplemental Tables S4 and S5). We also observed that LVLDLP levels may have a small causal effect on the *Distant* trait (IVW β = 0.008, p = 0.022) without any indication of pleiotropy or heterogeneity in the genetic instrument (Supplemental Tables S4 and S5). We observed significant horizontal pleiotropy between MVLDLC and *Avoidance* (MR-Egger intercept = - 0.001, p = 0.006), which appears to underlie a causal effect between them (MR-Egger β = 0.019, p = 3×10^−4^). To verify that the horizontal pleiotropy did not affect the causal estimate, we conducted a leave-one-out analysis to identify the pleiotropic variants from the MVLDLC genetic instrument (Supplemental Figure S2). When the pleiotropic variants from the genetic instrument were removed (MR-Egger intercept = −5×10^−4^, p = 0.21), the causal effect of MVLDLC levels on *Avoidance* was confirmed (IVW β = 0.007, p = 0.01; MR-Egger β = 0.013, p = 0.017; Weighted median β = 0.011, p = 0.012) (Figure 2b).

### Enrichment analysis

We performed a GO enrichment analysis based on the genetic instruments with significant causal effects on trauma-related traits to understand the biological processes involved. After FDR multiple testing correction (Q < 5%), we found several GO terms associated with Citrate levels (N=24), GP (N=28), LVLDLP (N=5) and MVLDLC (N=4) with respect to transcriptomic data from blood, but no enrichment based on brain transcriptomic information survived multiple testing correction (Supplemental Tables S6 and S7). Among the significant enrichments observed with respect to the genetic instrument tested, we verified a large similarity network that included multiple GO terms related to biological pathways implicated in brain development (Figure 3). Additionally, four GO terms were significant across GP, LVLDLP, and MVLDLC: GO:0001694~*histamine biosynthetic process*; GO:0004398~*histidine decarboxylase activity*; GO:0006547~*histidine metabolic process*; and GO:0042423~*catecholamine biosynthetic process*. Additionally, GO:0021954~*central nervous system neuron development* was significant in both GP and MVLDLC analyses (respectively, p = 2.29×10^−5^ and p = 3.73×10^−4^). The GP genetic instrument also showed enrichments for multiple GO terms related to brain function and regulation: GO:2000178~*negative regulation of neural precursor cell proliferation* (p = 3.08×10^−7^); GO:0097154~*GABAergic neuron differentiation* (p = 7.63×10^−4^); GO:2000977~*regulation of forebrain neuron differentiation* (p = 7.63×10^−4^); GO:0021533~*cell differentiation in hindbrain* (p = 1.37×10^−3^); GO:0021514~*ventral spinal cord interneuron differentiation* (p = 1.68×10^−3^); and GO:0021542~*dentate gyrus development* (p = 1.98×10^−3^). No brain enrichment or overlap to other analysis was observed with respect to the citrate genetic instrument, which showed enrichments with several metabolic processes with the most significant results observed for: GO:0015137~*citrate transmembrane transporter activity* (p = 2.88×10^−4^) and GO:0015746~*citrate transport* (p = 2.88×10^−4^).

**Figure 3:**
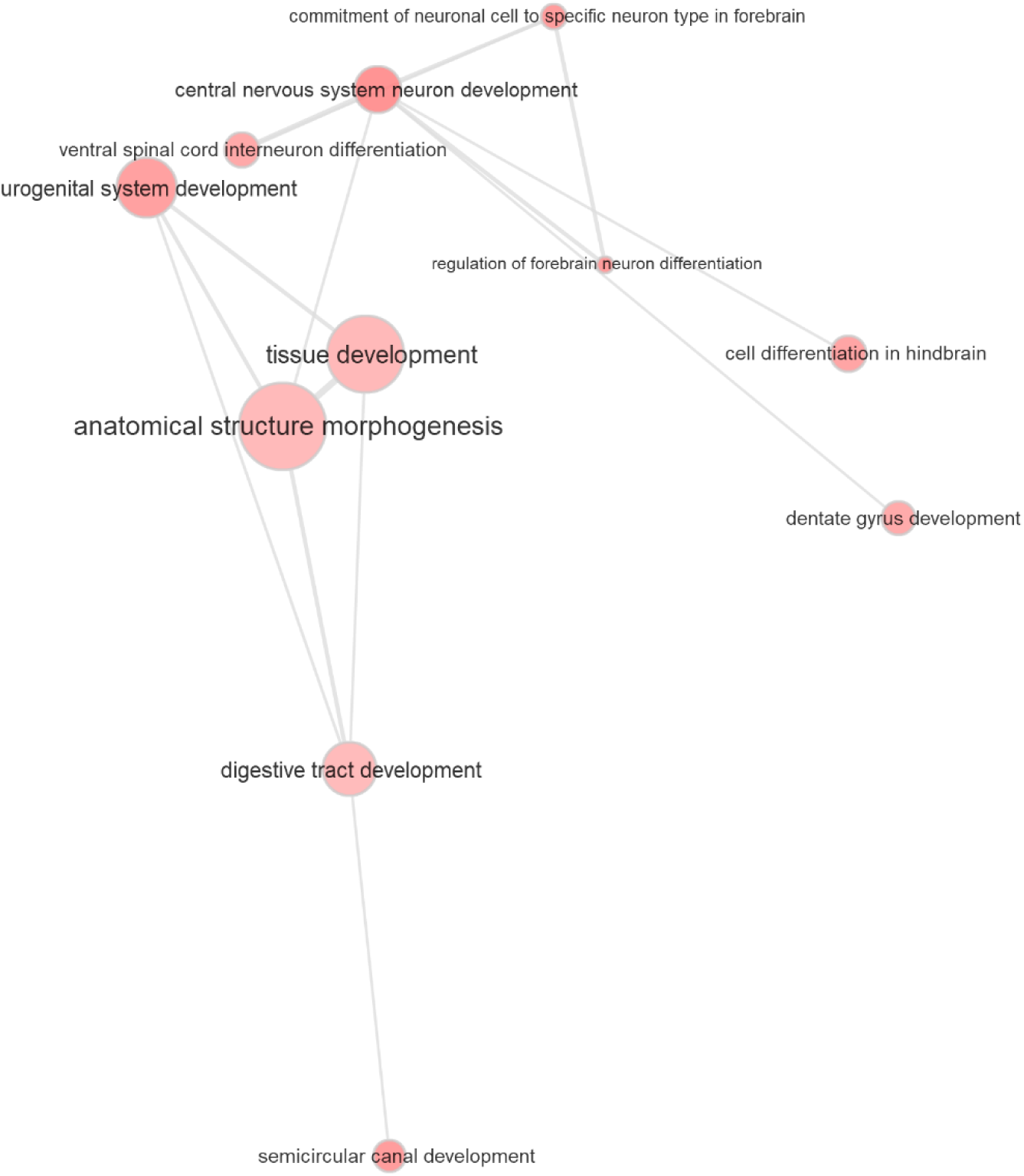
Similarity network based on GO enrichment results. Bubble color shading is proportional to the GO enrichment; bubble size is proportional to the frequency of the GO term in the Gene Ontology Annotation (UniProt) database. Edges in the graph represent the high similarity among the GO terms evaluated and the line width indicates the degree of similarity.

## Discussion

In the first study of its kind, our findings indicate that blood metabolite levels may be causally related to trauma response traits. Conversely, we did not find evidence of causal effects of trauma response traits on the metabolite levels investigated. These results reinforce the idea that metabolites may play a role in the pathophysiology of emotional and behavioral responses (23), and physical conditions (51) that occur after psychological trauma. In particular, we found evidence of causal effects with respect to four blood metabolites - citrate, GP, and two subclasses of VLDL (LVLDLP and MVLDLC).

Citrate is an important component of tricarboxylic acid (TCA). It is a substrate in the cellular energy metabolism cycle involved in the fatty acid synthesis, glycolysis, and gluconeogenesis (52). Furthermore, citrate is a chelating agent for divalent cations (e.g., Ca^2+^, Zn^2+^, and Mg^2+^), regulating these ions and potentially influencing the excitability of neurons acting, for example, via activity of glutamate receptors and NMDA receptors (52). Citrate has been linked with numerous biological processes, such as neurotransmitter synthesis and release, inflammation, insulin secretion, and histone acetylation (52–54). Low plasma citrate level was reported to be related to depression, suggesting that a reduction of TCA cycle substrates is associated with depressive symptoms (55). Our current analysis supports the causal relationship between low citrate levels and increased feelings of upset in response to traumatic events. This could be related to the role of citrate in several brain processes.

GP are acute-phase proteins, which are mainly represented by α-1-acid glycoprotein (AGP), also called orosomucoid (56). Increased GP plasma levels have been associated with immuno-modulating effects, drug-transporting proteins (e.g., histamine, serotonin), regulation of metabolism, and neuroinflammation (56, 57). AGP expression in astrocytes also appears to mediate astrocyte-microglia interactions during neuroinflammation (58, 59). In addition, GPs have been investigated as potential biomarkers for depression due to their ability to bind major classes of antidepressant drugs, such as tricyclic antidepressants and selective serotonin reuptake inhibitors (60). Furthermore, a recent study suggested that GPs were significantly associated with increased risk of dementia and lower general cognitive ability (26), which has in turn been correlated with risk for PTSD (61). These results support our findings suggesting an effect of glycoproteins on *Avoidance*.

VLDLs are categorized according to their molecular size (62). We identified two subclasses of VLDL (LVLDLP and MVLDLC) that may have causal effects on traits related to trauma response. LVLDLP and MVLDLC are lipoprotein particle subclasses of triglyceride-containing VLDL (62). Higher levels of VLDLs have been related to cardiovascular disorders (63) and type 2 diabetes (64), which are associated with traumatic stress and psychiatric disorders (65, 66). Elevated lipid levels have been previously observed in individuals affected by PTSD (67), but it is still unclear whether this relationship is due to potential confounding factors. Our findings suggest that two subclasses of VLDLs influence the trauma response, but further investigations will be needed to fully understand the molecular mechanisms involved.

The direction of causality between metabolites and trauma response was confirmed by enrichment analysis. Furthermore, the enrichment analysis showed that important biological processes for neuropsychiatric disorders were enriched in the genetic instruments related to the blood metabolites tested: histidine process (68), histamine (68), catecholamines (69) and citrate transport (52).

Previous studies have indicated that an exacerbated response to traumatic events is associated with both physiological and psychological dysfunction (9, 11). The response to emotional trauma is a complex phenomenon which triggers different regulatory systems in the brain and the body, with alterations in the hypothalamic- pituitary- adrenal axis (HPA) axis, neuroinflammatory processes and sympathetic nervous system, as well as hormonal and neurotransmitter responses (10, 11, 70). Our enrichment results suggest that biological processes linked with these metabolites (i.e., citrate, GP, LVLDLP, and MVLDLC) may be involved in the mechanism of trauma response.

Several previous studies have investigated the role of metabolites on trauma- and stressor-related disorders (71), but none investigated the same metabolites evaluated here. Different metabolites and glycans released from glycoproteins may be associated with mechanisms underlying PTSD (71). Our data suggest that some metabolites may be related to PTSD-related symptoms, for example, *Avoidance*.

In conclusion, this study is the first MR analysis to evaluate the causal relationship between metabolites and traits related to trauma response using genome-wide data. Our findings provide evidence that some metabolites (i.e., citrate, GP, and two VLDL subclasses) may have causal effects on emotional and behavioral responses to trauma: that is, they may modify mechanisms involved in the stress response. Future studies examining more metabolites and larger samples will be needed to dissect the molecular mechanisms by which blood metabolites affect brain regulation and function in determining the inter-individual variability in stress responses.

## Acknowledgments

We thank study participants, research groups, and the members of the cited studies for making their data available.

