## Supplemental Material for "Metabolome-Wide Mendelian Randomization Analysis of Emotional and Behavioral Responses to Traumatic Stress"

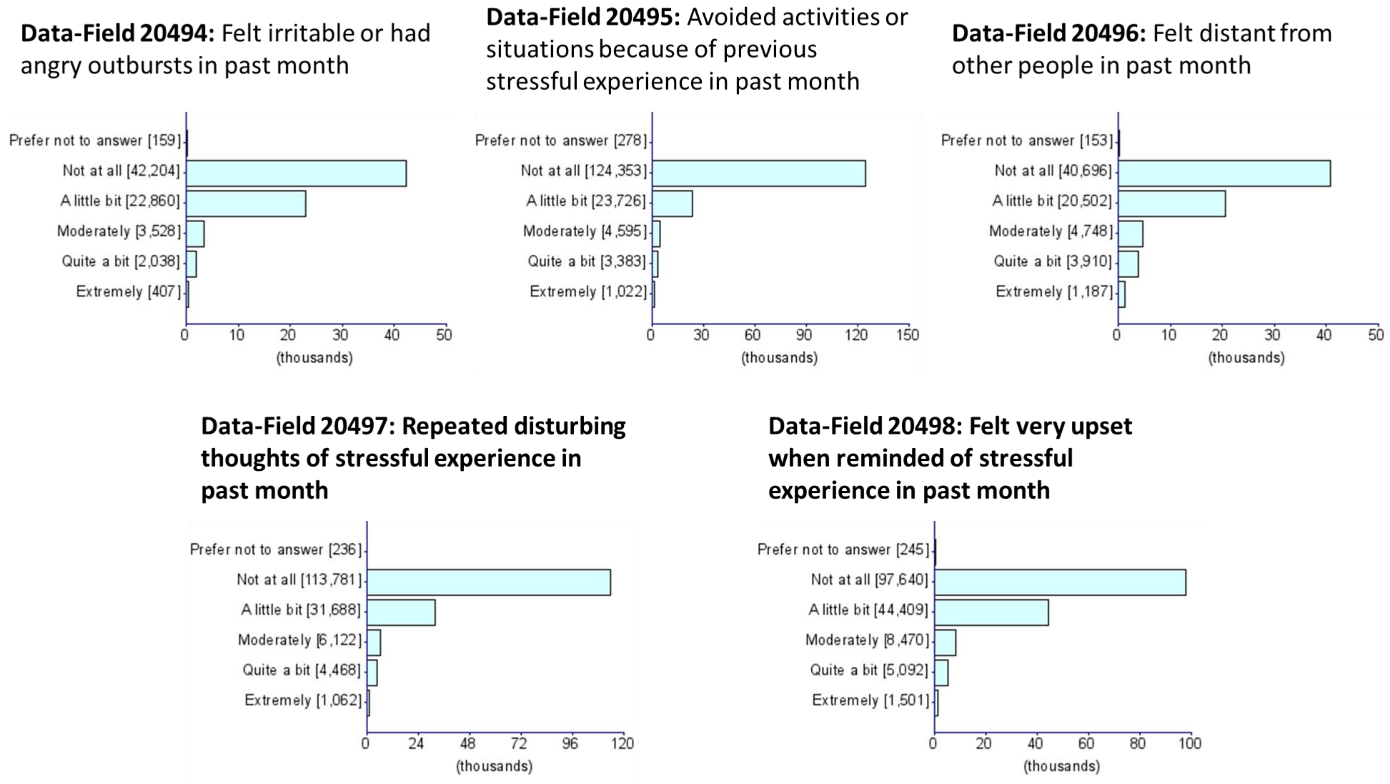


**Figure S1:** Distribution of the UKB participants' answers to the trauma-response questions.

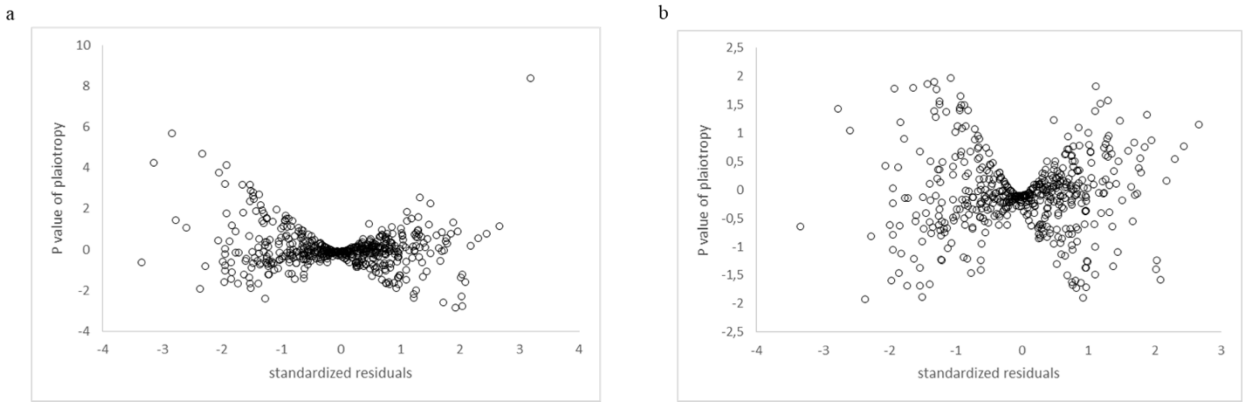


**Figure S2:** Leave-one-out analysis to identify potential outliers in total cholesterol in medium VLDL (MVLDLC) phenotype. **a.** Distribution of variants before leave-one-out analysis (N=544). **b.** Distribution of variants after removal of outliers (N=517).

**Table S1**: Heritability z-score of metabolites.

| Metabolites | Heritability | SE | Heritability z-score | p-value |
|---|---|---|---|---|
| 18:2 linoleic acid (LA) | 0.126 | 0.048 | 2.627 | 0.009 |
| 22:6 docosahexaenoic acid (DHA) | 0.130 | 0.037 | 3.481 | 4.99E-04 |
| 3Lhydroxybutyrate | 0.034 | 0.020 | 1.680 | 0.093 |
| Acetate | 0.053 | 0.020 | 2.670 | 0.008 |
| Acetoacetate | 0.076 | 0.028 | 2.736 | 0.006 |
| Alanine | 0.089 | 0.027 | 3.313 | 0.001 |
| Albumin | 0.060 | 0.025 | 2.447 | 0.014 |
| Apoprotein A1 | 0.078 | 0.031 | 2.495 | 0.013 |
| Apoprotein B | 0.082 | 0.039 | 2.089 | 0.037 |
| CH2 groups in fatty acids | 0.197 | 0.080 | 2.448 | 0.014 |
| CH2 groups to double bonds ratio | 0.057 | 0.043 | 1.336 | 0.182 |
| Cholesterol esters in large HDL (high-density lipoprotein) | 0.122 | 0.032 | 3.873 | 1.07E-04 |
| Cholesterol esters in large LDL (low-density lipoprotein) | 0.075 | 0.051 | 1.477 | 0.140 |
| Cholesterol esters in large VLDL (very-low density lipoprotein) | 0.163 | 0.034 | 4.749 | 2.04E-06 |
| Cholesterol esters in medium HDL | 0.052 | 0.029 | 1.819 | 0.069 |
| Cholesterol esters in medium LDL | 0.079 | 0.050 | 1.597 | 0.110 |
| Cholesterol esters in medium VLDL | 0.149 | 0.038 | 3.971 | 7.17E-05 |
| Cholesterol esters in very large HDL | 0.040 | 0.027 | 1.472 | 0.141 |
| Citrate | 0.073 | 0.021 | 3.441 | 0.001 |
| Concentration of chylomicrons and extremely large VLDL particles | 0.118 | 0.026 | 4.494 | 6.98E-06 |
| Concentration of IDL particles (intermediate-density lipoprotein) | 0.074 | 0.043 | 1.732 | 0.083 |
| Concentration of large HDL particles | 0.132 | 0.031 | 4.245 | 2.18E-05 |
| Concentration of large LDL particles | 0.074 | 0.050 | 1.480 | 0.139 |
| Concentration of large VLDL particles | 0.133 | 0.034 | 3.852 | 1.17E-04 |
| Concentration of medium HDL particles | 0.064 | 0.026 | 2.427 | 0.015 |
| Concentration of medium LDL particles | 0.078 | 0.049 | 1.610 | 0.107 |
| Concentration of medium VLDL particles | 0.151 | 0.035 | 4.251 | 2.12E-05 |
| Concentration of small HDL particles | 0.048 | 0.027 | 1.770 | 0.077 |
| Concentration of small LDL particles | 0.093 | 0.043 | 2.146 | 0.032 |
| Concentration of small VLDL particles | 0.148 | 0.037 | 3.968 | 7.25E-05 |
| Concentration of very large HDL particles | 0.063 | 0.027 | 2.324 | 0.020 |
| Concentration of very large VLDL particles | 0.127 | 0.030 | 4.292 | 1.77E-05 |
| Concentration of very small VLDL particles | 0.114 | 0.036 | 3.162 | 0.002 |
| Creatinine | 0.112 | 0.025 | 4.553 | 5.29E-06 |
| Double bonds in fatty acids | 0.166 | 0.054 | 3.053 | 0.002 |
| Esterified cholesterol | 0.062 | 0.051 | 1.211 | 0.226 |

| | | | | |
|---|---|---|---|---|
| Fatty acid length | 0.101 | 0.034 | 2.953 | 0.003 |
| Free cholesterol | 0.067 | 0.043 | 1.560 | 0.119 |
| Free cholesterol in IDL | 0.070 | 0.041 | 1.722 | 0.085 |
| Free cholesterol in large HDL | 0.106 | 0.029 | 3.676 | 2.37E-04 |
| Free cholesterol in large LDL | 0.063 | 0.048 | 1.308 | 0.191 |
| Free cholesterol in large VLDL | 0.124 | 0.030 | 4.154 | 3.27E-05 |
| Free cholesterol in medium HDL | 0.068 | 0.023 | 2.923 | 0.003 |
| Free cholesterol in medium VLDL | 0.110 | 0.034 | 3.296 | 0.001 |
| Free cholesterol in small VLDL | 0.110 | 0.033 | 3.304 | 0.001 |
| Free cholesterol in very large HDL | 0.049 | 0.025 | 1.925 | 0.054 |
| Glucose | 0.089 | 0.021 | 4.155 | 3.25E-05 |
| Glutamine | 0.063 | 0.021 | 2.976 | 0.003 |
| Glycerol | 0.031 | 0.021 | 1.473 | 0.141 |
| Glycine | 0.207 | 0.113 | 1.838 | 0.066 |
| Glycoproteins | 0.098 | 0.029 | 3.383 | 0.001 |
| HDL diameter | 0.121 | 0.030 | 4.057 | 4.97E-05 |
| Histidine | 0.032 | 0.024 | 1.367 | 0.172 |
| Isoleucine | 0.073 | 0.024 | 3.084 | 0.002 |
| Lactate | 0.022 | 0.018 | 1.201 | 0.230 |
| LDL diameter | 0.048 | 0.027 | 1.815 | 0.070 |
| Leucine | 0.048 | 0.021 | 2.250 | 0.024 |
| MonoLunsaturated fatty acids | 0.109 | 0.043 | 2.549 | 0.011 |
| Omega 3 fatty acids | 0.156 | 0.043 | 3.668 | 2.44E-04 |
| Omega 6 fatty acids | 0.068 | 0.046 | 1.485 | 0.138 |
| Omega 7 and 9 and saturated fatty acids | 0.092 | 0.044 | 2.112 | 0.035 |
| Other polyunsaturated fatty acids than 18:2 | 0.162 | 0.100 | 1.631 | 0.103 |
| Phenylalanine | 0.053 | 0.023 | 2.333 | 0.020 |
| Phosphatidylcholine and other cholines | NA | NA | NA | NA |
| Phospholipids in chylomicrons and extremely large VLDL | 0.097 | 0.027 | 3.614 | 3.01E-04 |
| Phospholipids in IDL | 0.061 | 0.041 | 1.472 | 0.141 |
| Phospholipids in large HDL | 0.126 | 0.031 | 4.104 | 4.06E-05 |
| Phospholipids in large LDL | 0.062 | 0.044 | 1.392 | 0.164 |
| Phospholipids in large VLDL | 0.116 | 0.031 | 3.745 | 1.80E-04 |
| Phospholipids in medium HDL | 0.067 | 0.023 | 2.880 | 0.004 |
| Phospholipids in medium LDL | 0.072 | 0.042 | 1.709 | 0.087 |
| Phospholipids in medium VLDL | 0.109 | 0.033 | 3.275 | 0.001 |
| Phospholipids in small VLDL | 0.115 | 0.035 | 3.290 | 0.001 |
| Phospholipids in very large HDL | 0.091 | 0.030 | 3.023 | 0.003 |
| Phospholipids in very large VLDL | 0.108 | 0.028 | 3.853 | 1.17E-04 |

| | | | | |
|---|---|---|---|---|
| Phospholipids in very small VLDL | 0.084 | 0.038 | 2.229 | 0.026 |
| Pyruvate | 0.036 | 0.021 | 1.742 | 0.082 |
| Ratio of bisLallylic bonds to double bonds in lipids | 0.294 | 0.111 | 2.663 | 0.008 |
| Ratio of bisLallylic bonds to total fatty acids in lipids | 0.270 | 0.099 | 2.744 | 0.006 |
| Serum total cholesterol | 0.064 | 0.038 | 1.671 | 0.095 |
| Serum total triglycerides | 0.124 | 0.036 | 3.453 | 0.001 |
| Sphingomyelins | -0.0004 | 0.04 | -0.01 | 0.992 |
| Total cholesterol in HDL | 0.094 | 0.029 | 3.291 | 0.001 |
| Total cholesterol in IDL | 0.070 | 0.045 | 1.536 | 0.124 |
| Total cholesterol in large HDL | 0.105 | 0.028 | 3.706 | 2.11E-04 |
| Total cholesterol in large LDL | 0.064 | 0.047 | 1.378 | 0.168 |
| Total cholesterol in large VLDL | 0.117 | 0.029 | 4.063 | 4.85E-05 |
| Total cholesterol in LDL | 0.068 | 0.050 | 1.339 | 0.180 |
| Total cholesterol in medium HDL | 0.053 | 0.025 | 2.087 | 0.037 |
| Total cholesterol in medium LDL | 0.068 | 0.047 | 1.455 | 0.146 |
| Total cholesterol in medium VLDL | 0.123 | 0.035 | 3.500 | 4.65E-04 |
| Total cholesterol in small LDL | 0.069 | 0.044 | 1.585 | 0.113 |
| Total cholesterol in small VLDL | 0.088 | 0.031 | 2.818 | 0.005 |
| Total cholesterol in very large HDL | 0.034 | 0.023 | 1.485 | 0.138 |
| Total fatty acids | 0.089 | 0.046 | 1.926 | 0.054 |
| Total lipids in chylomicrons and extremely large VLDL | 0.120 | 0.028 | 4.371 | 1.24E-05 |
| Total lipids in IDL | 0.072 | 0.044 | 1.640 | 0.101 |
| Total lipids in large HDL | 0.130 | 0.031 | 4.127 | 3.67E-05 |
| Total lipids in large LDL | 0.074 | 0.051 | 1.439 | 0.150 |
| Total lipids in large VLDL | 0.139 | 0.030 | 4.596 | 4.31E-06 |
| Total lipids in medium HDL | 0.060 | 0.026 | 2.258 | 0.024 |
| Total lipids in medium LDL | 0.080 | 0.050 | 1.606 | 0.108 |
| Total lipids in medium VLDL | 0.139 | 0.035 | 4.029 | 5.60E-05 |
| Total lipids in small HDL | 0.055 | 0.026 | 2.119 | 0.034 |
| Total lipids in small LDL | 0.086 | 0.045 | 1.892 | 0.059 |
| Total lipids in small VLDL | 0.142 | 0.038 | 3.792 | 1.49E-04 |
| Total lipids in very large HDL | 0.059 | 0.028 | 2.156 | 0.031 |
| Total lipids in very large VLDL | 0.138 | 0.030 | 4.568 | 4.92E-06 |
| Total lipids in very small VLDL | 0.104 | 0.035 | 2.913 | 0.004 |
| Total phosphoglycerides | 0.013 | 0.040 | 0.312 | 0.755 |
| Triglycerides in chylomicrons and extremely large VLDL | 0.092 | 0.027 | 3.374 | 0.001 |
| Triglycerides in IDL | 0.105 | 0.035 | 3.041 | 0.002 |
| Triglycerides in large VLDL | 0.113 | 0.029 | 3.886 | 1.02E-04 |
| Triglycerides in medium VLDL | 0.096 | 0.030 | 3.145 | 0.002 |

| | | | | |
|---|---|---|---|---|
| Triglycerides in small HDL | 0.063 | 0.027 | 2.324 | 0.020 |
| Triglycerides in small VLDL | 0.120 | 0.035 | 3.417 | 0.001 |
| Triglycerides in very large HDL | 0.084 | 0.025 | 3.291 | 0.001 |
| Triglycerides in very large VLDL | 0.122 | 0.030 | 4.108 | 3.99E-05 |
| Triglycerides in very small VLDL | 0.136 | 0.037 | 3.693 | 2.22E-04 |
| Tyrosine | 0.076 | 0.027 | 2.777 | 0.005 |
| Urea | 0.027 | 0.022 | 1.247 | 0.213 |
| Valine | 0.061 | 0.019 | 3.165 | 0.002 |
| VLDL diameter | 0.134 | 0.035 | 3.878 | 1.05E-04 |

*SE:* standard error.

**Table S2:** Genetic correlation among the traits related to trauma response.

| Metabolites | Heritability | rg | Standard error | p-value |
|---|---|---|---|---|
| Irritable | Upset | 0.7562 | 0.1233 | 8.58E-10 |
| Irritable | Repeated Thoughts | 0.8734 | 0.1233 | 1.38E-12 |
| Irritable | Distant | 0.4931 | 0.1751 | 0.0049 |
| Irritable | Avoidance | 0.5947 | 0.1335 | 8.40E-06 |
| Avoidance | Distant | 0.8241 | 0.1095 | 5.22E-14 |
| Avoidance | Repeated Thoughts | 0.9169 | 0.0439 | 7.72E-97 |
| Avoidance | Upset | 0.8828 | 0.0433 | 1.96E-92 |
| Distant | Repeated Thoughts | 0.8824 | 0.1181 | 7.73E-14 |
| Distant | Upset | 0.7652 | 0.098 | 5.77E-15 |
| Repeated Thoughts | Upset | 0.9398 | 0.0258 | 1.50E-290 |

***rg*: genetic correlation**

**Table S3:** Mendelian randomization (MR) results between metabolites (exposure) and trauma response phenotypes from the UK Biobank. Consistent effect directions and significant p-values are highlighted in bold.

| Exposure | Outcome | Method | N_SNP | Beta | SE | p-value |
|---|---|---|---|---|---|---|
| Citrate | Upset | MR-Egger | 8 | **-0.112** | 0.123 | 0.398 |
| | | Weighted median | 8 | **-0.046** | 0.023 | **0.048** |
| | | IVW | 8 | **-0.058** | 0.017 | **0.00091** |
| | | Simple mode | 8 | **-0.044** | 0.035 | 0.257 |
| | | Weighted mode | 8 | **-0.043** | 0.031 | 0.191 |
| Glycoproteins | Avoidance | MR-Egger | 488 | **0.004** | 0.006 | 0.511 |
| | | Weighted median | 488 | **0.007** | 0.004 | 0.127 |
| | | IVW | 488 | **0.008** | 0.003 | **0.003** |
| | | Simple mode | 488 | **0.0009** | 0.014 | 0.944 |
| | | Weighted mode | 488 | **0.0009** | 0.011 | 0.929 |
| | Repeated Thoughts | MR-Egger | 488 | **0.005** | 0.006 | 0.401 |
| | | Weighted median | 488 | **0.005** | 0.005 | 0.278 |
| | | IVW | 488 | **0.004** | 0.003 | 0.199 |
| | | Simple mode | 488 | **0.006** | 0.016 | 0.693 |
| | | Weighted mode | 488 | **0.006** | 0.012 | 0.603 |
| LVLDLP | Distant | MR-Egger | 2 747 | **0.011** | 0.006 | 0.137 |
| | | Weighted median | 2 747 | **0.009** | 0.006 | 0.132 |
| | | IVW | 2 747 | **0.008** | 0.003 | **0.022** |
| | | Simple mode | 2 747 | **0.201** | 0.452 | 0.655 |
| | | Weighted mode | 2 747 | **0.201** | 0.457 | 0.658 |
| MVLDLC | Avoidance | MR-Egger | 504 | **0.013** | 0.005 | **0.017** |
| | | Weighted median | 504 | **0.011** | 0.004 | **0.012** |
| | | IVW | 504 | **0.007** | 0.002 | **0.011** |
| | | Simple mode | 504 | **0.023** | 0.022 | 0.288 |
| | | Weighted mode | 504 | **0.016** | 0.017 | 0.357 |

*LVLDLP*: Concentration of large VLDL particles (Very-low-density lipoprotein). *MVLDLC*: Total cholesterol in medium VLDL. *N_SNP:* number of single nucleotide polymorphism evaluated. *SE:* standard error. *IVW:* inverse variance weighted.

**Table S4:** Pleiotropy tests for variants included in Mendelian randomization (MR) analysis.

| Causality tests | MR-Egger intercept | SE | p-value |
|---|---|---|---|
| CIT→ Upset | 0.004 | 0.01 | 0.67 |
| GP→Avoidance | 0.0004 | 0.0004 | 0.365 |
| GP→Repeated Thoughts | -0.00013 | 0.0005 | 0.814 |
| LVLDLP→Distant | -0.00016 | 0.0005 | 0.744 |

*SE:* standard error. *LVLDLP:* Concentration of large VLDL particles (Very-low-density lipoprotein).

**Table S5:** Heterogeneity tests for variants included in Mendelian randomization (MR) analysis.

| Causality tests | Methods | Q-test | Q-pvalue |
|---|---|---|---|
| CIT→ Upset | MR-Egger | 4.82 | 0.56 |
| | IVW | 5.01 | 0.65 |
| GP→Avoidance | MR-Egger | 504.11 | 0.275 |
| | IVW | 504.97 | 0.277 |
| GP→Repeated Thoughts | MR-Egger | 534.01 | 0.065 |
| | IVW | 534.06 | 0.069 |
| LVLDLP→Distant | MR-Egger | 2593.78 | 0.98 |
| | IVW | 2593.89 | 0.981 |

*CIT:* citrate. *GP:* glycoproteins acetyls. *LVLDLP:* Concentration of large VLDL particles (Very-low-density lipoprotein). *IVW:* inverse variance weighted.

**Table S6:** Significant gene ontology (GO) molecular pathways enriched in blood tissue.

| Metabolite | GO_ID | GO_Term | p-value | Q-value |
|---|---|---|---|---|
| Glycoproteins | GO:2000178 | negative regulation of neural precursor cell proliferation | 3.08E-07 | 6.16E-05 |
| | GO:0001694 | histamine biosynthetic process | 5.09E-05 | 1.70E-03 |
| | GO:0004398 | histidine decarboxylase activity | 5.09E-05 | 1.70E-03 |
| | GO:0006547 | histidine metabolic process | 5.09E-05 | 1.70E-03 |
| | GO:0021954 | central nervous system neuron development | 2.29E-05 | 1.70E-03 |
| | GO:0042423 | catecholamine biosynthetic process | 5.09E-05 | 1.70E-03 |
| | GO:0021902 | commitment of neuronal cell to specific neuron type in forebrain | 7.63E-04 | 1.31E-02 |
| | GO:0035854 | eosinophil fate commitment | 7.63E-04 | 1.31E-02 |
| | GO:0060872 | semicircular canal development | 7.63E-04 | 1.31E-02 |
| | GO:0070345 | negative regulation of fat cell proliferation | 7.88E-04 | 1.31E-02 |
| | GO:0097154 | GABAergic neuron differentiation | 7.63E-04 | 1.31E-02 |
| | GO:2000977 | regulation of forebrain neuron differentiation | 7.63E-04 | 1.31E-02 |
| | GO:0001655 | urogenital system development | 9.66E-04 | 1.49E-02 |
| | GO:0043434 | response to peptide hormone | 1.12E-03 | 1.60E-02 |
| | GO:0021533 | cell differentiation in hindbrain | 1.37E-03 | 1.83E-02 |
| | GO:0021514 | ventral spinal cord interneuron differentiation | 1.68E-03 | 2.10E-02 |
| | GO:0021542 | dentate gyrus development | 1.98E-03 | 2.33E-02 |
| | GO:0008532 | N-acetyllactosaminide beta-1,3-N-acetylglucosaminyltransferase activity | 2.36E-03 | 2.46E-02 |
| | GO:0030311 | poly-N-acetyllactosamine biosynthetic process | 2.29E-03 | 2.46E-02 |
| | GO:0045650 | negative regulation of macrophage differentiation | 2.46E-03 | 2.46E-02 |
| | GO:0006707 | cholesterol catabolic process | 3.50E-03 | 3.34E-02 |
| | GO:0070742 | C2H2 zinc finger domain binding | 4.29E-03 | 3.90E-02 |
| | GO:0031369 | translation initiation factor binding | 5.61E-03 | 4.49E-02 |
| | GO:0032057 | negative regulation of translational initiation in response to stress | 5.48E-03 | 4.49E-02 |
| | GO:0045654 | positive regulation of megakaryocyte differentiation | 5.35E-03 | 4.49E-02 |
| | GO:0009888 | tissue development | 5.89E-03 | 4.53E-02 |
| | GO:0048565 | digestive tract development | 6.47E-03 | 4.79E-02 |
| | GO:0043303 | mast cell degranulation | 6.97E-03 | 4.98E-02 |
| LVLDLP | GO:0001694 | histamine biosynthetic process | 1.98E-04 | 2.69E-02 |
| | GO:0004398 | histidine decarboxylase activity | 1.98E-04 | 2.69E-02 |
| | GO:0006547 | histidine metabolic process | 1.98E-04 | 2.69E-02 |
| | GO:0042423 | catecholamine biosynthetic process | 1.98E-04 | 2.69E-02 |
| | GO:0021954 | central nervous system neuron development | 3.73E-04 | 4.05E-02 |
| MVLDLC | GO:0001694 | histamine biosynthetic process | 1.07E-04 | 1.18E-02 |
| | GO:0004398 | histidine decarboxylase activity | 1.07E-04 | 1.18E-02 |
| | GO:0006547 | histidine metabolic process | 1.07E-04 | 1.18E-02 |
| | GO:0042423 | catecholamine biosynthetic process | 1.07E-04 | 1.18E-02 |
| Citrate | GO:0015137 | citrate transmembrane transporter activity | 2.88E-04 | 4.76E-03 |
| | GO:0015746 | citrate transport | 2.88E-04 | 4.76E-03 |

| | | | | |
|---|---|---|---|---|
| Citrate | GO:0030135 | coated vesicle | 4.75E-04 | 5.22E-03 |
| | GO:0030130 | clathrin coat of trans-Golgi network vesicle | 7.52E-04 | 5.48E-03 |
| | GO:0097443 | sorting endosome | 8.31E-04 | 5.48E-03 |
| | GO:0046326 | positive regulation of glucose import | 1.28E-03 | 7.04E-03 |
| | GO:0006094 | gluconeogenesis | 3.09E-03 | 1.45E-02 |
| | GO:0005905 | coated pit | 6.01E-03 | 1.73E-02 |
| | GO:0009653 | anatomical structure morphogenesis | 6.28E-03 | 1.73E-02 |
| | GO:0030136 | clathrin-coated vesicle | 5.17E-03 | 1.73E-02 |
| | GO:0035338 | long-chain fatty-acyl-CoA biosynthetic process | 4.82E-03 | 1.73E-02 |
| | GO:0042147 | retrograde transport, endosome to Golgi | 5.98E-03 | 1.73E-02 |
| | GO:0019432 | triglyceride biosynthetic process | 7.06E-03 | 1.79E-02 |
| | GO:0005770 | late endosome | 1.16E-02 | 2.15E-02 |
| | GO:0005802 | trans-Golgi network | 1.11E-02 | 2.15E-02 |
| | GO:0005819 | spindle | 1.05E-02 | 2.15E-02 |
| | GO:0006006 | glucose metabolic process | 1.17E-02 | 2.15E-02 |
| | GO:0006898 | receptor-mediated endocytosis | 1.12E-02 | 2.15E-02 |
| | GO:0005198 | structural molecule activity | 1.40E-02 | 2.44E-02 |
| | GO:0007067 | mitotic nuclear division | 1.95E-02 | 3.22E-02 |
| | GO:0006886 | intracellular protein transport | 2.43E-02 | 3.65E-02 |
| | GO:0044255 | cellular lipid metabolic process | 2.38E-02 | 3.65E-02 |
| | GO:0004871 | signal transducer activity | 2.56E-02 | 3.68E-02 |
| | GO:0005975 | carbohydrate metabolic process | 3.31E-02 | 4.55E-02 |

*GO_ID:* gene ontology identification. *GO_Term:* name of pathway

**Table S7:** Top 50 molecular pathways for concentration of large very-low-density lipoproteins particles (LVLDP) from enrichment analyses of brain tissue.

| GO_ID | GO_Term | p-value | Q-value |
|---|---|---|---|
| GO:0030126 | COPI vesicle coat | 0.0015 | 0.0792 |
| GO:0043234 | protein complex | 0.0030 | 0.0792 |
| GO:0001525 | angiogenesis | 0.0031 | 0.0792 |
| GO:0001077 | transcriptional activator activity, RNA polymerase II core promoter proximal region sequence-specific binding | 0.0033 | 0.0792 |
| GO:0003682 | chromatin binding | 0.0036 | 0.0792 |
| GO:0000978 | RNA polymerase II core promoter proximal region sequence-specific DNA binding | 0.0042 | 0.0792 |
| GO:0006366 | transcription from RNA polymerase II promoter | 0.0068 | 0.0792 |
| GO:0060038 | cardiac muscle cell proliferation | 0.0070 | 0.0792 |
| GO:0003700 | transcription factor activity, sequence-specific DNA binding | 0.0096 | 0.0792 |
| GO:0045944 | positive regulation of transcription from RNA polymerase II promoter | 0.0144 | 0.0792 |
| GO:0006355 | regulation of transcription, DNA-templated | 0.0202 | 0.0792 |
| GO:0046534 | positive regulation of photoreceptor cell differentiation | 0.0235 | 0.0792 |
| GO:0035257 | nuclear hormone receptor binding | 0.0266 | 0.0792 |
| GO:0043522 | leucine zipper domain binding | 0.0275 | 0.0792 |
| GO:0008048 | calcium sensitive guanylate cyclase activator activity | 0.0276 | 0.0792 |
| GO:0005307 | Choline: sodium symporter activity | 0.0281 | 0.0792 |
| GO:0008292 | acetylcholine biosynthetic process | 0.0281 | 0.0792 |
| GO:0015220 | choline transmembrane transporter activity | 0.0281 | 0.0792 |
| GO:0015871 | choline transport | 0.0282 | 0.0792 |
| GO:0031284 | positive regulation of guanylate cyclase activity | 0.0282 | 0.0792 |
| GO:0033265 | choline binding | 0.0287 | 0.0792 |
| GO:0022400 | regulation of rhodopsin mediated signaling pathway | 0.0289 | 0.0792 |
| GO:0005615 | extracellular space | 0.0295 | 0.0792 |
| GO:0097381 | photoreceptor disc membrane | 0.0295 | 0.0792 |

| | | | |
|---|---|---|---|
| GO:0001523 | retinoid metabolic process | 0.0303 | 0.0792 |
| GO:0050896 | response to stimulus | 0.0313 | 0.0792 |
| GO:0042572 | retinol metabolic process | 0.0318 | 0.0792 |
| GO:0016056 | rhodopsin mediated signaling pathway | 0.0347 | 0.0792 |
| GO:0005070 | SH3/SH2 adaptor activity | 0.0351 | 0.0792 |
| GO:0007274 | neuromuscular synaptic transmission | 0.0356 | 0.0792 |
| GO:0007622 | rhythmic behavior | 0.0356 | 0.0792 |
| GO:0021660 | rhombomere 3 formation | 0.0356 | 0.0792 |
| GO:0021666 | rhombomere 5 formation | 0.0356 | 0.0792 |
| GO:0021612 | facial nerve structural organization | 0.0357 | 0.0792 |
| GO:0035284 | brain segmentation | 0.0357 | 0.0792 |
| GO:0004666 | prostaglandin-endoperoxide synthase activity | 0.0358 | 0.0792 |
| GO:0032227 | negative regulation of synaptic transmission, dopaminergic | 0.0358 | 0.0792 |
| GO:0042633 | hair cycle | 0.0359 | 0.0792 |
| GO:0050473 | arachidonate 15-lipoxygenase activity | 0.0359 | 0.0792 |
| GO:0071837 | HMG box domain binding | 0.0359 | 0.0792 |
| GO:0090050 | positive regulation of cell migration involved in sprouting angiogenesis | 0.0359 | 0.0792 |
| GO:0090271 | positive regulation of fibroblast growth factor production | 0.0359 | 0.0792 |
| GO:0009750 | response to fructose | 0.0360 | 0.0792 |
| GO:0042562 | hormone binding | 0.0361 | 0.0792 |
| GO:0090336 | positive regulation of brown fat cell differentiation | 0.0362 | 0.0792 |
| GO:0090362 | positive regulation of platelet-derived growth factor production | 0.0363 | 0.0792 |
| GO:0014037 | Schwann cell differentiation | 0.0364 | 0.0792 |
| GO:0010226 | response to lithium ion | 0.0366 | 0.0792 |
| GO:0031394 | positive regulation of prostaglandin biosynthetic process | 0.0366 | 0.0792 |
